# Isolation, screening, degradation characteristics of a quinclorac-degrading bacteria D and its potential of bioremediation for rice field environment polluted by quinclorac

**DOI:** 10.1101/2021.04.14.439927

**Authors:** Siqi Huang, Jiuyue Pan, Mancuo Tuwang, Hongyan Li, Chenyi Ma, Mingxue Chen, Xiaoyan Lin

## Abstract

Quinclorac (QNC) is a highly selective, hormonal, and low-toxic herbicide with a long duration. And the growth and development of subsequent crops are easily affected by QNC accumulated in the soil. In this paper, a QNC-degrading strain D was isolated and screened from the rice paddy soil. Through morphology, physiological and biochemical tests and 16Sr DNA gene analysis, strain D was identified as *Cellulosimicrobium cellulans* sp. And the QNC degradation characteristics of strain D were studied. Under the optimal culture conditions, the QNC-degrading rate was 45.9% after culturing for 21 days. The QNC-degrading efficiency of strain D in the field was evaluated by a simulated pot experiment. The results show that strain D can promote the growth of rice and QNC-degrading effectively. This research could provide a new bacterial species for microbial degradation of QNC and lay a theoretical foundation for further research on QNC remediation.

**Importance:** At present, some QNC-degrading bacteria have been isolated from different environments, but there are no reports of *Cellulosimicrobium cellulans* sp. bacterial that could degrade QNC. In this study, a new QNC-degradation strain was selected from the paddy soil. The degradation characteristics of strain D were studied in detail. The results shown that strain D had a satisfactory quinclorac-degrading efficiency. Two degradation products of QNC by strain D were identified by HPLC-Q-TOF/MS: 3-pyridylacetic acid (138.0548 m/z) and 3-ethylpyridine (108.0805 m/z), which have not been reported before. The strain D had a potential ability of quinclorac-degrading effectively in the quinclorac-polluted paddy field environment.

## Introduction

Quinclorac (QNC) is a highly selective hormone and low toxicity herbicide, which is mainly used to control barnyardgrass in rice fields (1). It was widely used because of its good herbicidal effect, wide application period, small application dosage and long-lasting time. According to “Southern Pesticides” report, QNC was one of the top 10 pesticides used in rice herbicides in China, and its annual saled can reach 41.21 million US dollars (2). However, due to the long-lasting usage of QNC, it is easy to accumulate in the soil and affect the growth and development of subsequent crops (3, 4). In Hunan Province, Jiangxi Province, Guangdong Province and other regions of China, the tobacco-rice rotation cropping model was often adopted. When the spray amount of QNC exceeding 25% of the recommended application dose in the rice season, it was harmful to subsequent crops, increased the deformity rate of tobacco leaves and reduced the yield of tobacco (5–7); the residual phytotoxicity of QNC also seriously threatened solanaceae crops, umbelliferae crops and chenopodiaceae crops. After the application of QNC, crops such as potatoes, peppers, carrots, celery, spinach, beets, etc. were sensitive to this herbicide. They would not be planted until at least two years later (8–10). Moreover, the residual QNC in the environment could affect the growth and development of animals adversely, caused abnormal development or reproductive abnormalities, and further affected the diversity and abundance of natural organisms (11). With the wide application of QNC in rice production, its harmed to animals and plants has attracted more and more attention. Therefore, it is necessary to find a practical and efficient method to degrade QNC residues and solve the environmental pollution problem caused by QNC residues.

The current degradation methods of QNC are mainly chemical degradation and biodegradation. Chemical methods mainly include photodegradation and electrodynamic soil cleaning. Photodegradation was achieved by using titanium dioxide to catalyze the photodegradation of QNC in paddy field water and ultrapure water. TiO_2_ P25 was used to degrade QNC completely in ultrapure water within 40 minutes under the irradiation of 1100 W xenon lamp, however, it took 130 minutes for QNC to be completely degraded in paddy water (12, 13). The degradation-products of photodegradation became more complex, and QNC was not yet fully mineralized. The light absorption range of TiO_2_ P25 is limited to the ultraviolet region of 320-400 nm, and ultraviolet light only accounts for 5% of the entire solar spectrum. Therefore, most of the experiments of photocatalytic degradation of pollutants were carried out under the irradiation of high-power xenon lamps or ultraviolet lamps. It required a lot of electrical energy, which limited large-scale experiments. Electrokinetic soil cleaning (14) is one of the most promising technologies for remediation of soil polluted with pesticides. However, during the electrochemical process, the initial pH value of the soil would change, which led to degradation of agricultural soil. In addition, during the electric soil cleaning process, pollutants were removed from the soil and then dissolved in the medium of water. Therefore, additional treatment was required to eliminate pollutants in the water, and this process was tedious and complicated. Due to the diverse metabolic pathways of microorganisms, extensive substrates and few by-products, microorganisms have broad application prospects in pesticide degradation.

At present, some QNC-degrading bacteria have been isolated from different environments. *Bordetella* sp. HN36 could degrade not only QNC, but also quinoline, phthalic acid, phenol and catechol (15). The pot experiment shown that *Alcaligenes* sp. J3 had obvious repairing effect on phytotoxic tobacco (16). Results of the bioremediation experiment of strain *Pantoea* QC06 shown that it also had a significant repair effect on tobacco phytotoxicity, and had certain practical applications for degrading QNC residues in tobacco fields (17). *Arthrobacter* sp. MC-10 could degrade QNC residues in contaminated soil within 7 days (18). Li Yingying et al. shown that the degradation product of this strain *Mycobacterium* sp. F4 was 3-chloro-7-hydroxyquinine-8-carboxylic acid or 7-chloro-3-hydroxyquinine-8-carboxylic acid, and the degradation products was non-phytotoxic to tobacco (19). Lang et al. screened out *Streptomyces* sp. AH-B through circulating fluidized bed culture and enrichment, the main degradation product were 3-chloro-7-methoxy-8-quinoline carboxylic acid, 3-chloro-7-methyl-8-Quinoline carboxylic acid, 3-chloro-7-oxyethyl-8-quinoline-carboxylic acid, and 3,7-dichloro-6-methyl-8-quinoline carboxylic acid (20). Research by Zhou Ting et al. found that the QNC-degrading efficiency of *Stenotrophomonas maltophilia* J03 in MSM medium was 33.5% after 21 days, and it could effectively alleviate the harm of QNC to tobacco plant growth (21). The QNC-degrading bacteria screened in most of the existing studies are used to remediate phytotoxic tobacco, and there were few studies on the remediation of QNC residual pollution in paddy soil. The purpose of this study is to screen out QNC-degrading bacteria adapted to the paddy field environment, with the hope that it will be widely used in QNC residue-contaminated paddy fields.

Until now, there are no reports of *Cellulosimicrobium cellulans* sp. bacterial that could degrade QNC. Current research shown that *Cellulosimicrobium cellulans* sp. bacterial could not only decompose cellulose, alkane powder, gelatin, xylan, and even paraffin, but also fixed nitrogen (22); Liang et al. isolated and screened out the *cellulosimicrobium cellulans* DGNK-JJ1, which could effectively dissolve potassium. Provided effective strains for the production of potassium-dissolving bacterial fertilizers and contributed to the development of ecological agriculture (23); Ferrer et al. shown that *Cellulosimicrobium cellulans* sp. bacterial could be used as biocontrol strains for plant diseases (24). These results revealed that this kind of microorganism could be used in fertilizers, plant protection and extraction of protective agents, etc., and had a excellent biological research prospect and development value. And this study found that *Cellulosimicrobium cellulans* sp. bacterial had a new characteristic of satisfactory QNC-degrading effect. Firstly, in this study, a strain of QNC-degrading bacteria D was isolated and screened from the paddy soil. It was identified as a *Cellulosimicrobium cellulans* sp. bacteria. Then through experiments, the degradation effect, degradation characteristics and degradation products of this strain were determined. Finally, the actual application of the strain was evaluated by pot simulation test.

## Results and discussion

### Identification of strain D

In our experiment, a strain that could degrade QNC effectively was obtained through isolation and screening, and named as strain D. The colony was round, umbilical, light yellow, moist, smooth surface and neat edges. The cell was rod-shaped, swollen cysts or spores, thicker capsule, chain-like arrangement, and gram positive. Its physiological and biochemical properties were shown in Table S1. The glucose, D-ribose, and contact enzyme tests are positive, and the raffinose test is negative. This result was consistent with the description of *Cellulosimicrobium cellulans* sp. bacterial in the “Berger’s Bacterial Identification Manual” (8th edition).

**Table S1.**
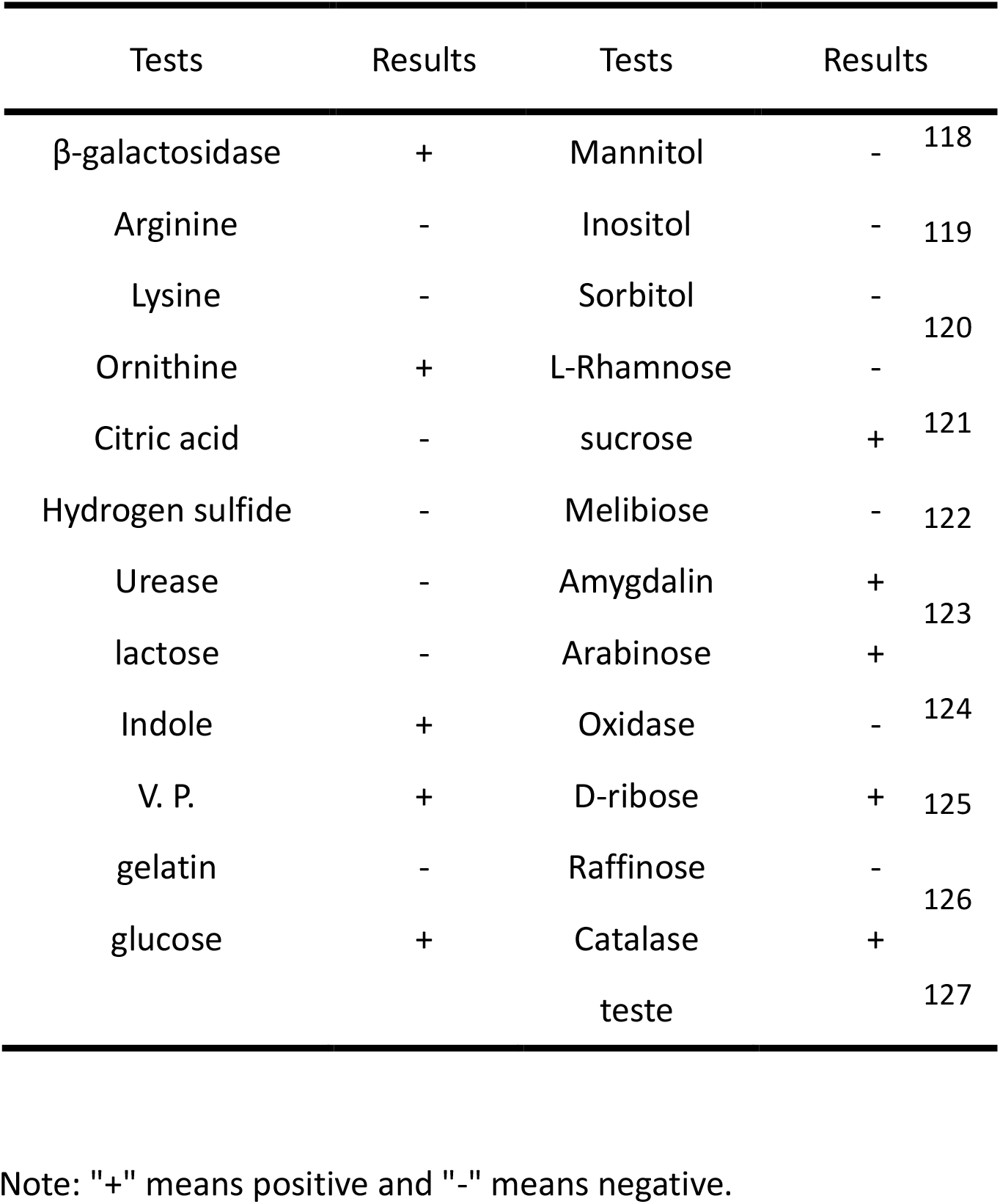
Physiological and biochemical characteristics of strain D

The 16S rRNA sequence of strain D was uploaded to NCBI for nucleotide sequence comparison, and constructed a phylogenetic tree. Fig. S2 is the 16S rDNA phylogenetic tree of strain D. It can be seen from the Fig. S2, strain D (1384 bp, GenBank accession no. MW404399) and *Cellulosimicrobium cellulans* DSM 43879 (GenBank accession no. NR_119095) are clustered in the same branch on the 16S rRNA phylogenetic tree, with homology is 99.93%. Therefore, strain D was identified as *Cellulosimicrobium cellulans* strain.

**Fig. S2.**
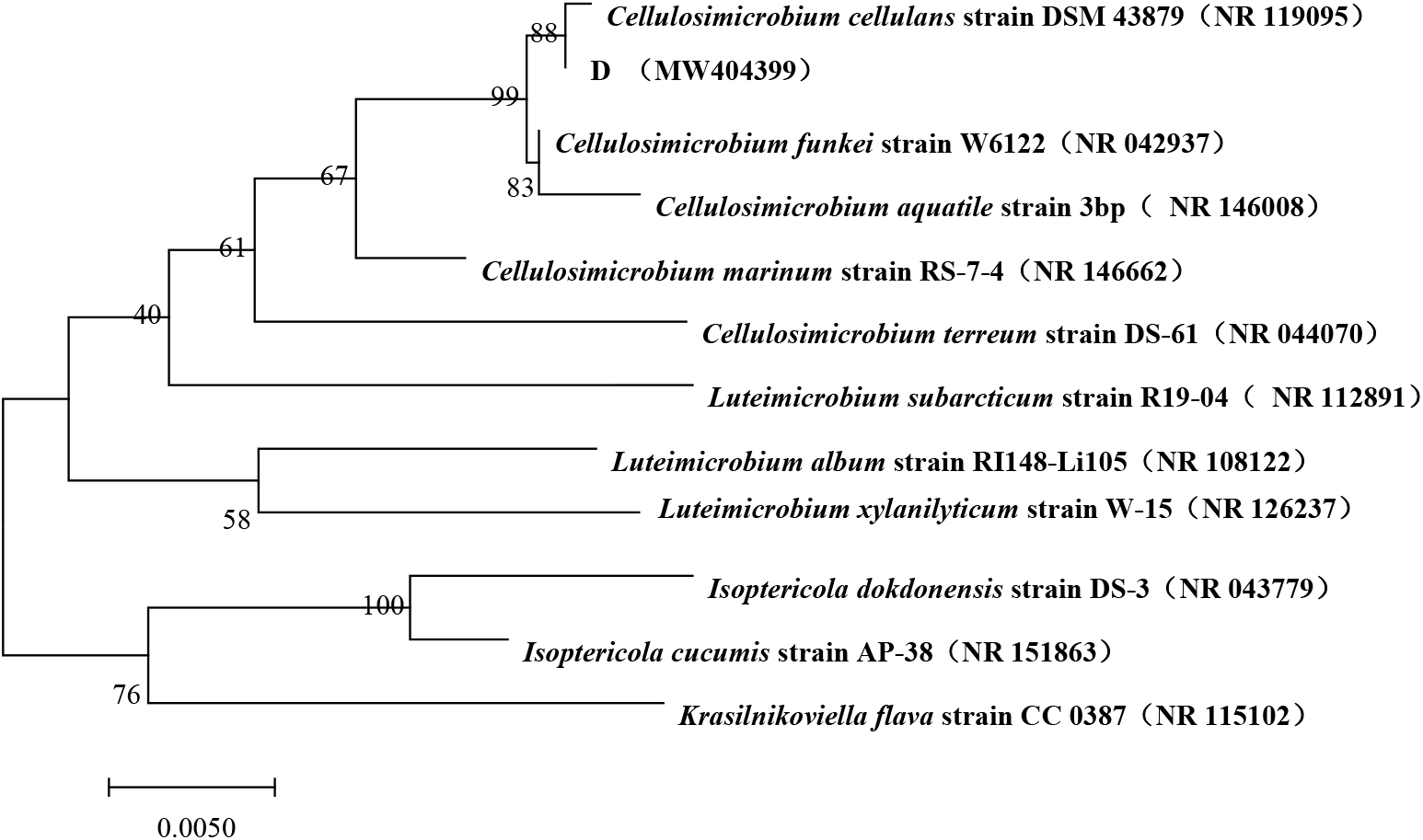
Phylogenetic tree of the 16S rDNA of strain D

### Degradation characteristics of QNC by strain D

#### The effect of initial pH on the degradation of QNC by strain D

It can be seen from Fig. S3(A) that there was a poor QNC-degrading effect by strain D in an environment with strong acidity (pH=4) and strong alkaline (pH=10). When the pH of the culture solution was 6, strain D had the best degradation effect in a weakly acidic environment, and the degradation rate was 31.6%. The chemical structure of QNC contains easily hydrolyzable carboxyl groups, with a pH of 4.35, which is weakly acidic (25). QNC is easy to dissociate in alkaline conditions, and acidic conditions can inhibit its dissociation. The suitable pH value for rice growth is 6-7, so QNC is not easy to dissociate naturally in the paddy soil. The strains selected in this study had a better degradation effect than other conditions when the pH value was 6-7. And they are very suitable for degrading QNC in rice fields under natural environment.

**Fig. S3.**
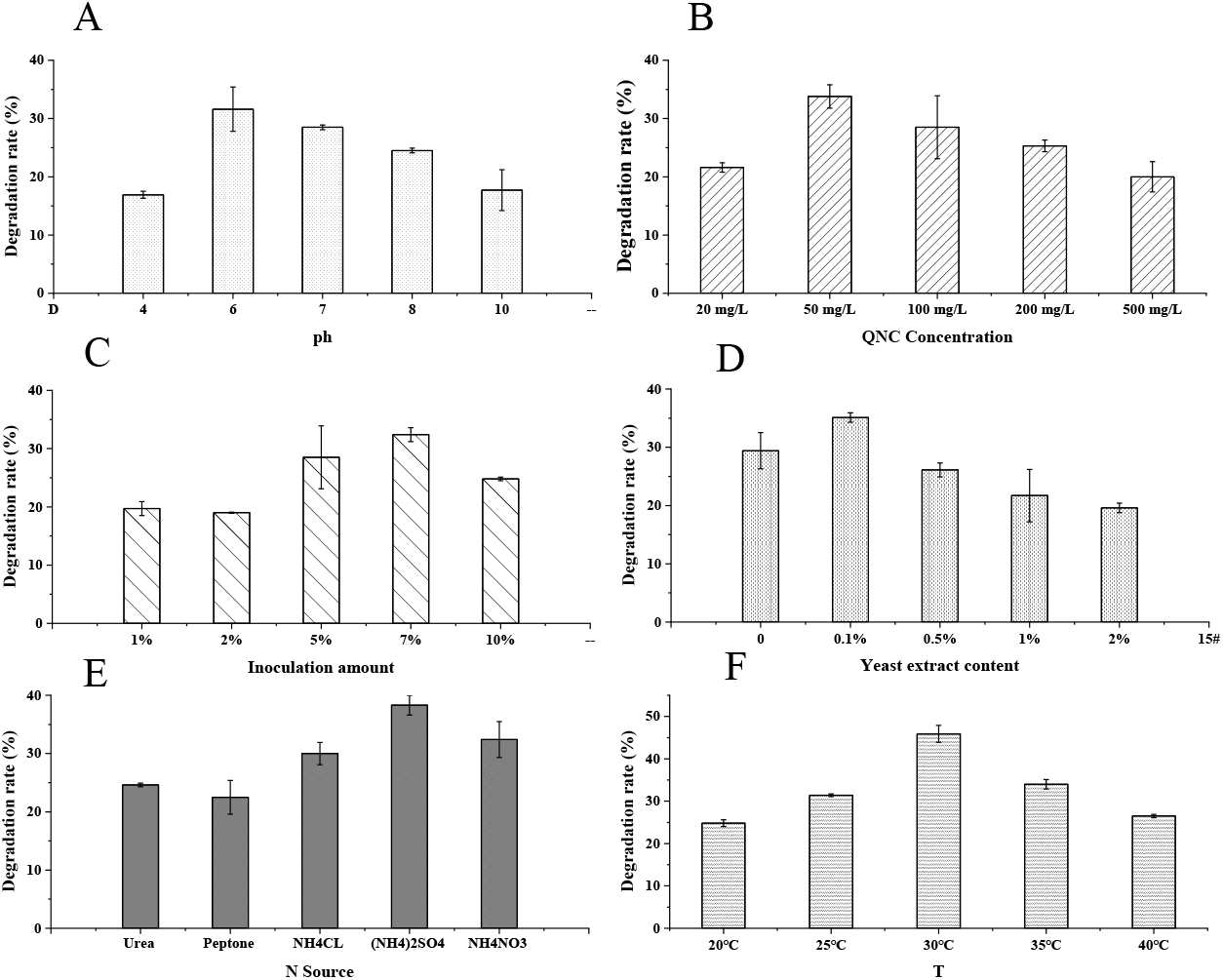
Factors affecting QNC degradation. A: Effect of initial pH on QNC-degradation by strain D; B: Effect of initial concentration of QNC on QNC-degradation by strain D; C: Effect of inoculation on the QNC-degradation effect by strain D; D: Effect of external nitrogen source on QNC-degradation by strain D; E: Effect of yeast extract content on the QNC-degradation effect by strain D; F: Effect of temperature on QNC-degradation by strain D.

### The effect of initial QNC on the degradation efficiency of QNC

The degradation effect of strain D in the MSM liquid medium with the initial concentration of QNC at 20, 50, 100, 200, 500 mg·L^−1^ was shown in Fig. S3(B). When the concentration of QNC was 50 mg·L^−1^, strain D had the best degradation effect, and the degradation rate was 33.8%. As the carbon source of the strain, QNC had a great influence on the degradation effect of strain D. Within a certain range of 20-50 mg·L^−1^, the degradation rate increased with the increase of QNC concentration. But when the concentration was too high, the degradation rate of QNC was at a lower level. The reason might be that the high concentration of physiological toxicity of QNC produces inhibited the reproduction and metabolism of the strain D. Studied by Lv Zhenmei (26) shown that QNC concentrations exceeding a certain limit would cause oxidative stress to microorganisms. The optimal initial mass concentration of QNC reported is in the range of 5-1000 mg·L^−1^. The *Stenotrophomonas maltophilia* J03 (21) 21d can degrade 33.5% of QNC at the initial mass concentration of 5 mg·L^−1.^. And the *Burkholderia cepacia* WZ1 (26) 5 d can degrade 90% of QNC. But degrading bacteria with low or high optimal QNC concentration are not suitable for common QNC contaminated farmland (375 g·hm^−2^·a^−1^). The optimal QNC concentration of strain D isolated by us was closer to the QNC concentration in seriously polluted paddy soil, which is suitable for heavily polluted paddy fields.

### The influence of the inoculum size on the degradation efficiency of QNC

The influence of the inoculation amount on the degradation efficiency of QNC was shown in Fig. S3(C). The growth of microorganisms would be affected by the amount of inoculum, which in turn affected the growth and reproduction of strains. If the amount of inoculation was little, the bacterial culture would have a longer delay time, and then as the amount of inoculation increased, the degradation rate of the strain would show an upward trend. Strain D had the best degradation-effect when the inoculation amount was 7%, and it could continuously and stably degrade QNC. The degradation rate of QNC could reach 32.4% after 21 days of culture. When the inoculum amount exceeded 7%, the degradation rate shown a downward trend, which may be due to the inhibition of competition between bacteria, resulting in insufficient nutrients and affecting the degradation rate of QNC.

### The effect of N source on the degradation efficiency of QNC

The type and content of nutrients affect the degradation ability of microorganisms. By optimizing the type of N source, the degradation efficiency of strain D could be effectively improved. The effect of different N sources on the degradation efficiency of QNC was shown in Fig. S3(D). It could be seen that the degradation of QNC was different when the culture medium contained different kinds of nitrogen sources. The promotion effect of different N sources on degradation of QNC by strain D was: (NH_4_)_2_SO_4_>NH_4_NO_3_>NH_4_Cl>Urea>Peptone. Therefore, (NH_4_)_2_SO_4_ was selected as the optimal N source. When using (NH_4_)_2_SO_4_ as the N source, strain D had the best QNC-degradation effect with the degradation rate of 38.3% in 21 days. When other conditions were the same, the nitrogen source was (NH_4_)_2_SO_4_, the strain degradation rate could be increased by 5.9% to 15.8%. Moreover, (NH_4_)_2_SO_4_ is an excellent nitrogen fertilizer, suitable for general soil and crops. It can make branches and leaves grow vigorously, improve fruit quality and yield, and enhance crop resistance to disasters. It can be used as base fertilizer, top dressing and seed fertilizer, widely used in rice production.

### The effect of yeast extract content on the degradation efficiency of QNC

Yeast extract is rich in protein, amino acids, peptides, nucleotides, B vitamins, and trace elements. It can not only be used as a supplement for N and C sources, but also contains vitamins and micronutrients that are beneficial to the growth of microorganisms (27). Appropriate addition of yeast extract could promote the growth and reproduction of microorganisms and improve the degradation efficiency. The effect of the content of yeast extract on the degradation of QNC by strain D was shown in Fig. S3(E). When the content of yeast extract was 0.1%, strain D had the best degradation effect, with a degradation rate of 35.1%. When yeast extract was not added and when the amount of yeast extract was too much, the QNC-degradation effect of strain D was not as good as the addition of 0.1% yeast extract. When the amount of yeast extract was 2%, the degradation effect of 21d was the worst. Perhaps the large amount of yeast extract changed the degradation-target of strain D.

### The effect of temperature on the degradation of QNC by strain D

Temperature is an important physical factor affecting the growth and survival of microorganisms. Temperature mainly affected the QNC-degrading effect of strain D by affecting the growth, reproduction and metabolism of microorganisms. The optimal temperature of QNC degrading bacteria had been reported to be between 25°C and 35°C In the same other culture conditions, at different temperatures, the degradation of QNC by strain D was shown in Fig. S3(F). It could be seen from the figure that as the culture temperature increased, the ability of strain D to degrade QNC gradually increased. When the culture temperature reached 30°C the degradation effect was the best, and the degradation rate was 45.9%. Then the degradation effect decreased with increasing temperature. Strain D had a degradation rate of 31.4%-45.9% in an environment between 25 °C and 35 °C which shown that the strain could adapt to the temperature during rice planting and growth.

### QNC Degradation of optimal culture conditions

In the optimal culture conditions, the initial pH was 6, the initial QNC concentration was 50 mg·L^−1^, the inoculation amount was 7%, the yeast extract content was 0.1%, and the nitrogen source was (NH_4_)_2_SO_4_, the culture temperature was 30°C and cultured for 21d. The degradation of QNC by strain D and the growth curve were shown in Fig. S4. Strain D could degrade 45.9% of QNC after 21 days of culture. Strain D entered the logarithmic growth phase after 3 days of culture, and entered the stable phase after 18 days. QNC could only be degraded by light and microorganisms under natural conditions. The QNC-degrading bacteria screened in most of the existing studies were used to repair phytotoxic tobacco, and there were few studies on the repair of QNC residual pollution in paddy soil (17, 28). The purpose of this study was to screen out QNC-degrading bacteria adapted to the paddy field, with the hope that it will be widely used in QNC residue-contaminated paddy fields. The degradation effect of QNC-degrading bacteria was related to many factors, including temperature, pH value, initial substrate mass concentration, inoculum amount, and the degradation ability of the strain, etc (29). In this study, a single factor experiment was used to optimize the degradation conditions of strain D. Finally, it was found that the degradation conditions of strain D were similar to the basic paddy field environment, and strain D could adapt to the paddy field environment.

**Fig. S4.**
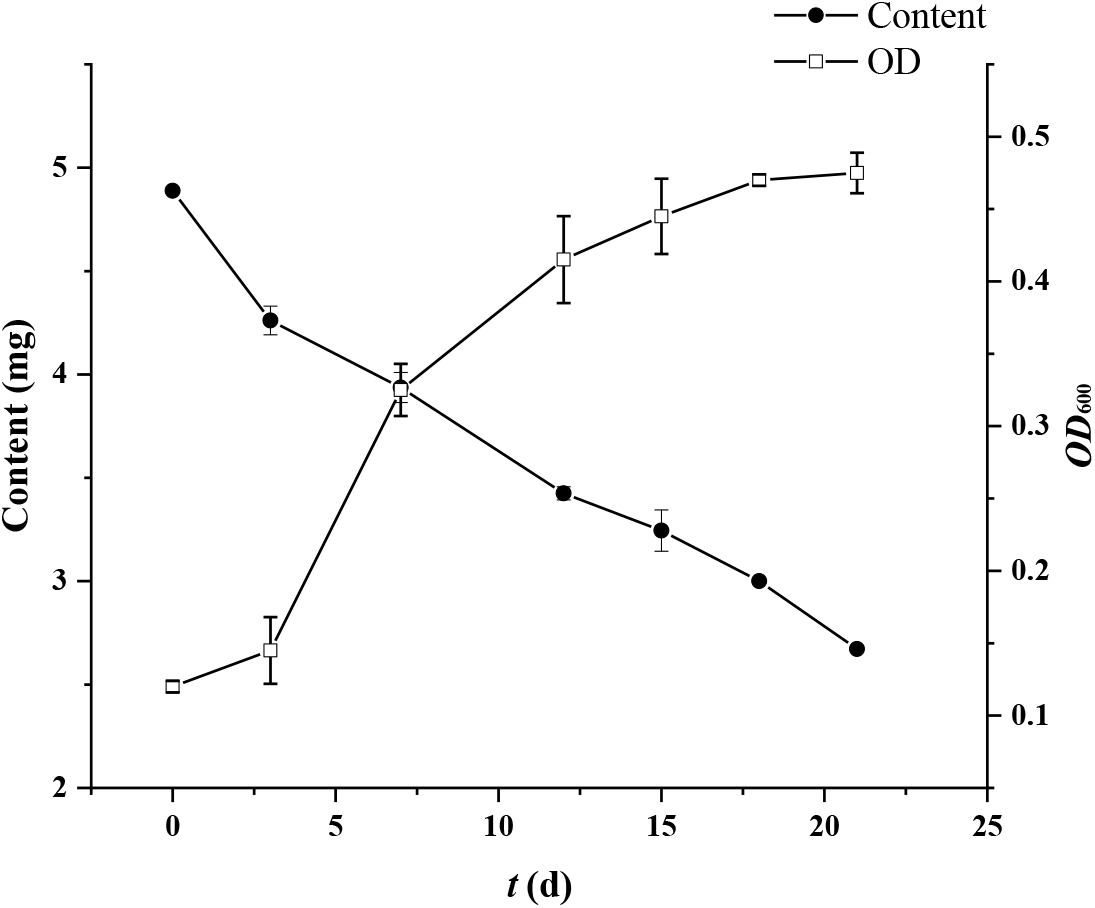
The effect of on the optimal conditions on the QNC-degradation of strain D and growth curve

### Analysis of degradation products of QNC by strain D

The QuEChERS method was used to extract QNC and its degradation products from MSM medium at different culture stages. QNC and its degradation products were detected by HPLC-Q-TOF/MS. Two possible QNC metabolites had been detected. The mass spectrum and chromatogram of the detected QNC and the two degradation products were shown in Fig. S5. The main degradation products confirmed by mass spectrometry were 3-Pyridylacetic acid (138.0548 m/z) and 3-Ethylpyridine, (108.0805 m/z). Due to the lack of standards, degradation products have not been quantified. These products had not been reported in previous studies on the QNC-degradation, and the degradation pathways and enzymes involved are still in study. These compounds were probably new metabolites of QNC degraded by the strain D. Of course, further tests were needed to verify our hypothesis.

**Fig. S5.**
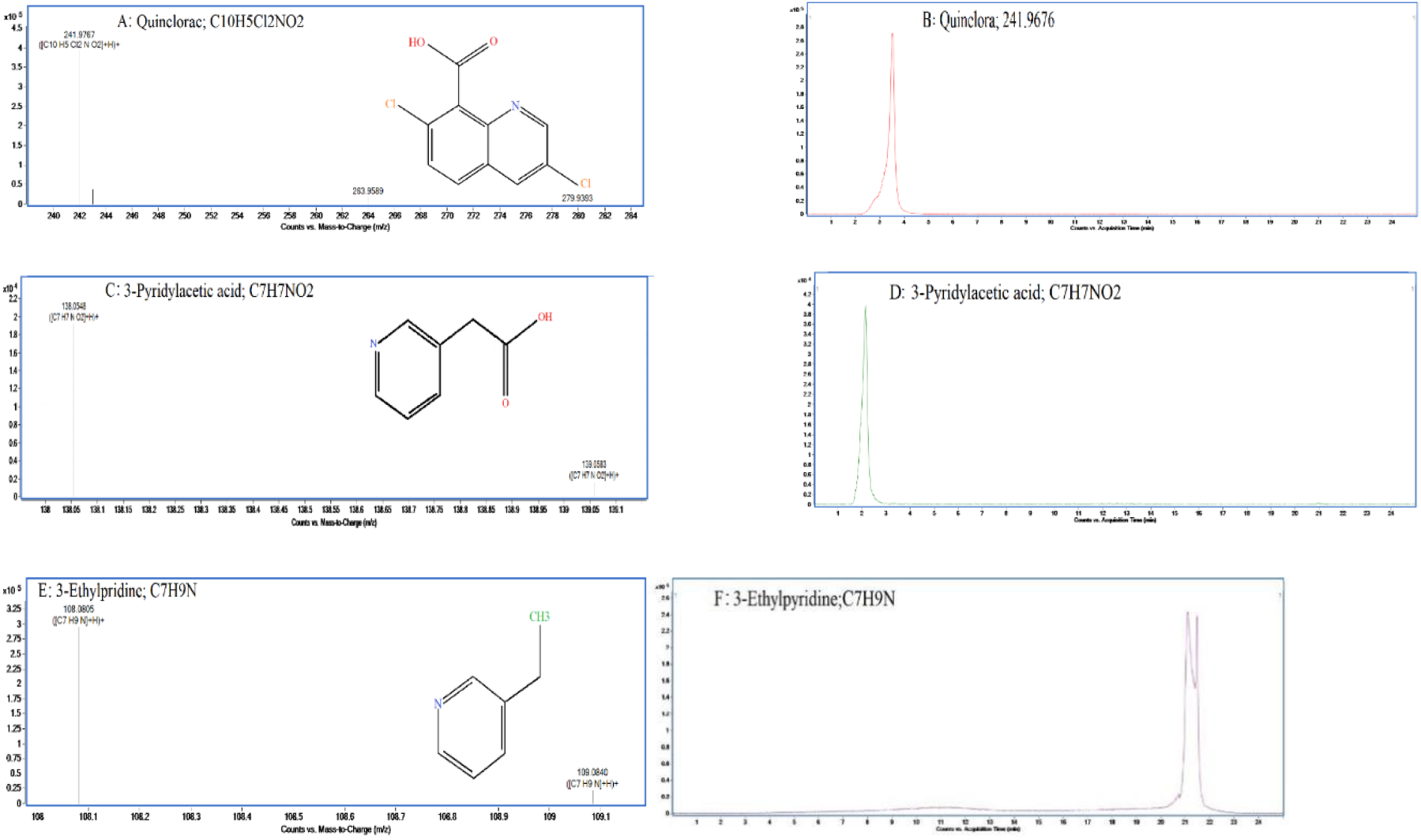
The mass spectrum and chromatogram of QNC and QNC degradation products. A: The mass spectrum of QNC; B: The mass chromatogram of QNC; C: The mass spectrum of 3-Pyridylacetic acid; D: The mass chromatogram of 3-Pyridylacetic acid; E: The mass spectrum of 3-Ethylpyridine; F: The mass chromatogram of 3-Ethylpyridine.

### The effect evaluation of strain D on the degradation of QNC in simulated rice fields conditions

In a potted simulated paddy field environment, the recommended dosage and twice the dosage were applied for soil restoration experiments. In this experiment, part of the soil was sterilized, and added QNC-degrading bacteria to explore the field bioremediation effect of QNC-polluted. In the 28-day pot experiment, the degradation of QNC in the soil in each treatment group was shown in Fig. S6. Experiments on normal soil shown (Fig. S6A and S6B) that strain D can effectively accelerate the degradation of QNC in normal soil. When the QNC concentration was 750 g·hm^−2^·a^−1^ in the environment, the degradation effect was better. Experiments on sterilized soil shown (Fig.S6C and S6D) that the degradation rate of QNC in the control group without degradation bacteria was significantly slower than that with degradation bacteria. Similarly, in an treatment with a QNC concentration of 750 g·hm^−2^·a^−1^, strain D had a stronger degradation-ability of degrading 75.45% of QNC within 21 days. It could be seen that the strain D was very suitable for the seriously polluted rice field environment. By comparing the degradation of QNC in the experimental group in the normal soil and the sterilized soil (Fig.S6E and S6F), it could be found clearly that QNC was degraded faster in the normal soil. There were two possible reasons, First, in the sterilized soil, only the degradation-bacteria D played a degradation-effect, while exogenous additive QNC-degrading strains in the normal soil also played a certain role in promoting the degradation of QNC. When exogenous additive QNC-degrading strains and degradation bacteria D had a synergistic effect, the degradation effect of QNC would be better. Second, after the soil was sterilized, its physical and chemical properties may be changed, and the degradation bacteria couldn’t perform better degradation in this condition. Whether the soil was sterilized or not, the degradation bacteria D could accelerate the degradation of QNC, which indicated that degradation bacteria D could adapt to the paddy field environment, effectively degraded QNC, and repaired QNC-contaminated rice fields.

**Fig. S6.**
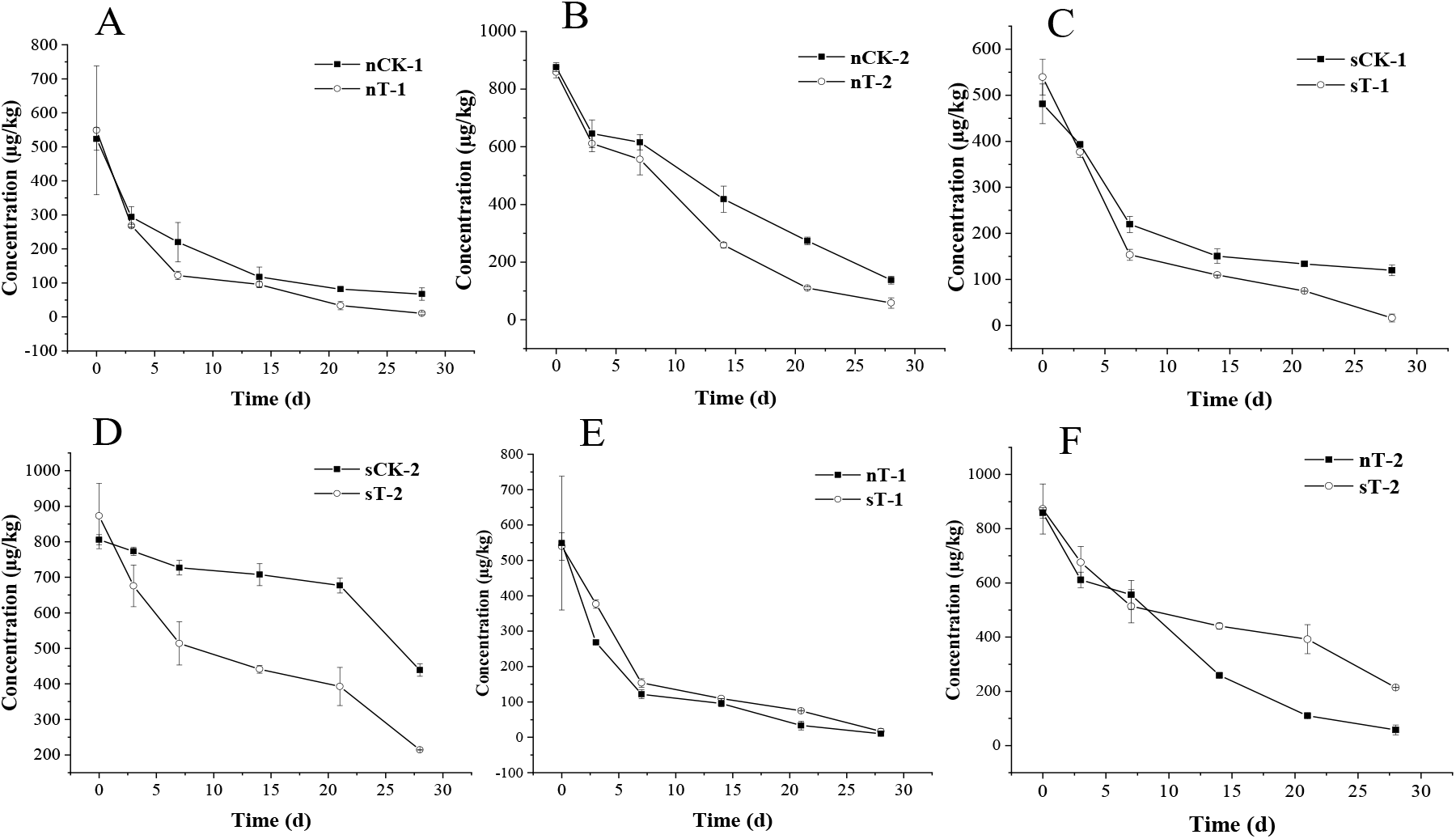
The degradation of QNC in the soil. A: In normal soil, the degradation of the recommended dosage; B: In normal soil, the degradation of 2 times the recommended dosage; C: In sterilized soil, the degradation of the recommended dosage; D: In sterilized soil, the degradation of 2 times the recommended dosage; E: Degradation of the experimental group with the recommended dosage in sterilized and normal soil; F: Degradation of the experimental group at 2 times the recommended dose in sterilized and normal soil.

### The effect evaluation of strain D on the growth of rice plants in simulated rice fields conditions

Sensitive dicotyledons may be significantly affected by high concentrations of QNC, such as chlorosis, necrosis, and growth inhibition (30). Although rice had a relatively high resistance to QNC, a higher risk would be brought to the growth of rice seedlings if QNC was used excessively. This study explored the effects of strain D on rice physiology under QNC stress through pot simulation experiments. In the pot simulation experiment, the application of degradation-bacteria had obvious effects on the plant. After spraying 7d, when the QNC concentration in the pot was at the recommended dose (375 g·hm^−2^·a^−1^) (Fig. S7A and S7B), the rice plant height of the normal experimental groups were 20.5% higher than that of the control group. The fresh stem weight of the sterilized experimental group was 64.3% heavier than that in the sterilized control group. When the QNC concentration in the pot was 2 times the recommended dose (750 g·hm^−2^·a^−1^) (Fig. S7C and S7D), only the sterilized and normal experimental groups had significant differences in the dry root weight of rice from the control group. There were obvious differences in the roots of plants, but not in other parts. The reason may be that rice root exudates interacted with rhizosphere soil microorganisms to promote root growth (31). After 21 days, when the QNC concentration in the pot was 2 times the recommended dose (750 g·hm^−2^·a^−1^) (Fig.8C and 8D), compared with the control group, the rice plant height, root length and stem fresh weight of the sterilized and normal experimental groups were significantly different after 21 days. However, this situation did not appear on the 7th day of the test. The reason might be that high concentrations of QNC had a greater impact on plant physiology, and QNC-degrading bacteria needed more time to adapt to the environment before they could begin to play a role. And the root dry weight of the normal experimental group was 41.4% heavier than that of the sterilized experimental group. This situation was consistent with the previous phenomenon of faster degradation in the normal experimental group. Through the pot experiment, it could be seen that the QNC-degrading bacteria D had a significant bioremediation effect on QNC-polluted soil. It showed that in the QNC-polluted paddy field environment, degrading bacteria D could promote the degradation of residual QNC and reduce its phytotoxicity to rice.

**Fig. S7.**
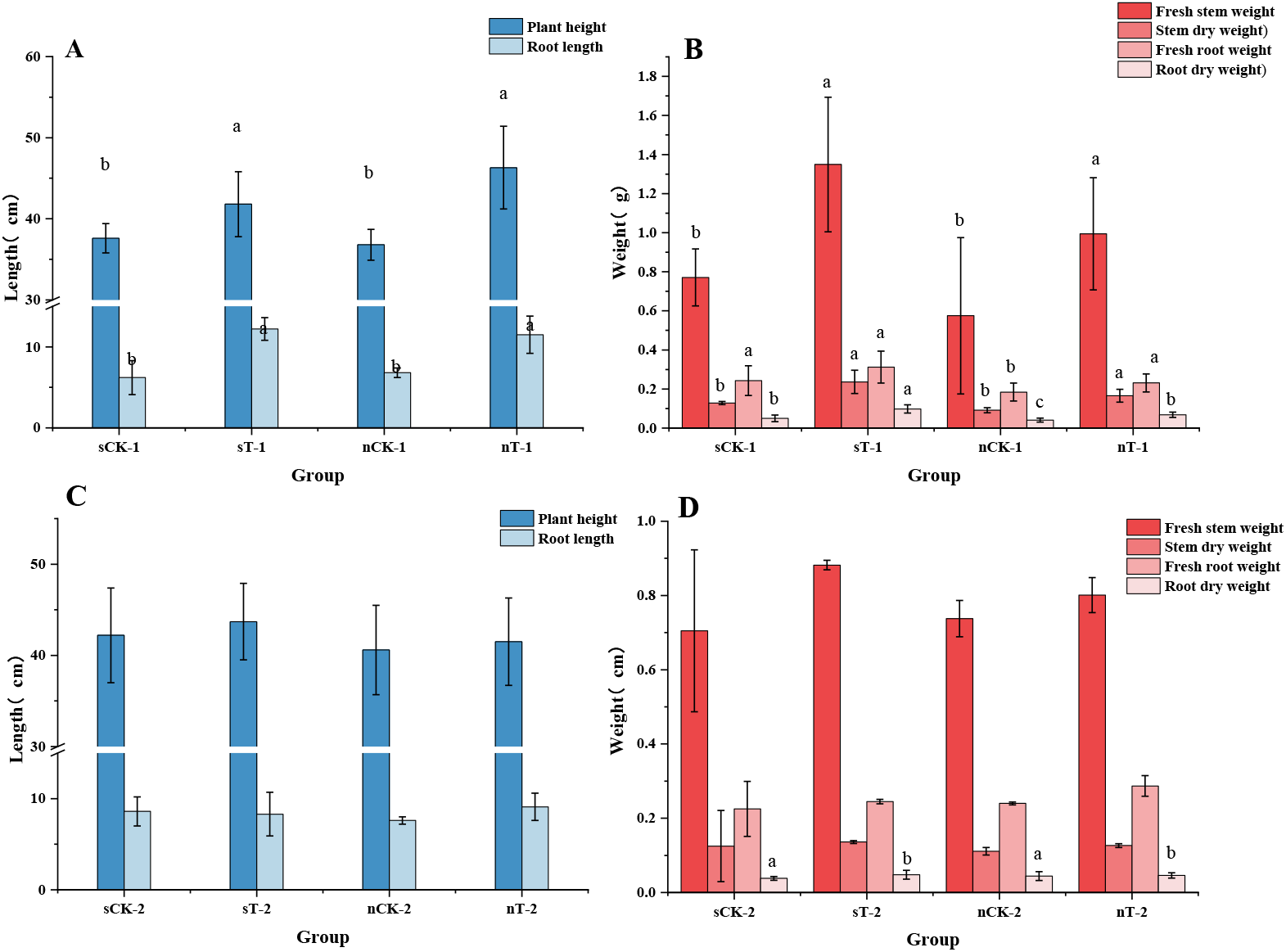
The growth of rice plants in simulated rice fields conditions in 7d. Note that different letters indicate significant differences at the 0.05 level. A : plant height and root length of experimental group and control group at recommended dose (375 g·hm^−2^·a^−1^); B: fresh stem weight, stem dry weight, fresh root weight and root dry weight of experimental group and control group at recommended dose (375 g·hm^−2^·a^−1^); C: plant height and root length of experimental group and control group at 2 times recommended dose (750 g·hm^−2^·a^−1^); D: fresh stem weight, stem dry weight, fresh root weight and root dry weight of experimental group and control group at 2 times recommended dose (750 g·hm^−2^·a^−1^).

**Fig. S8.**
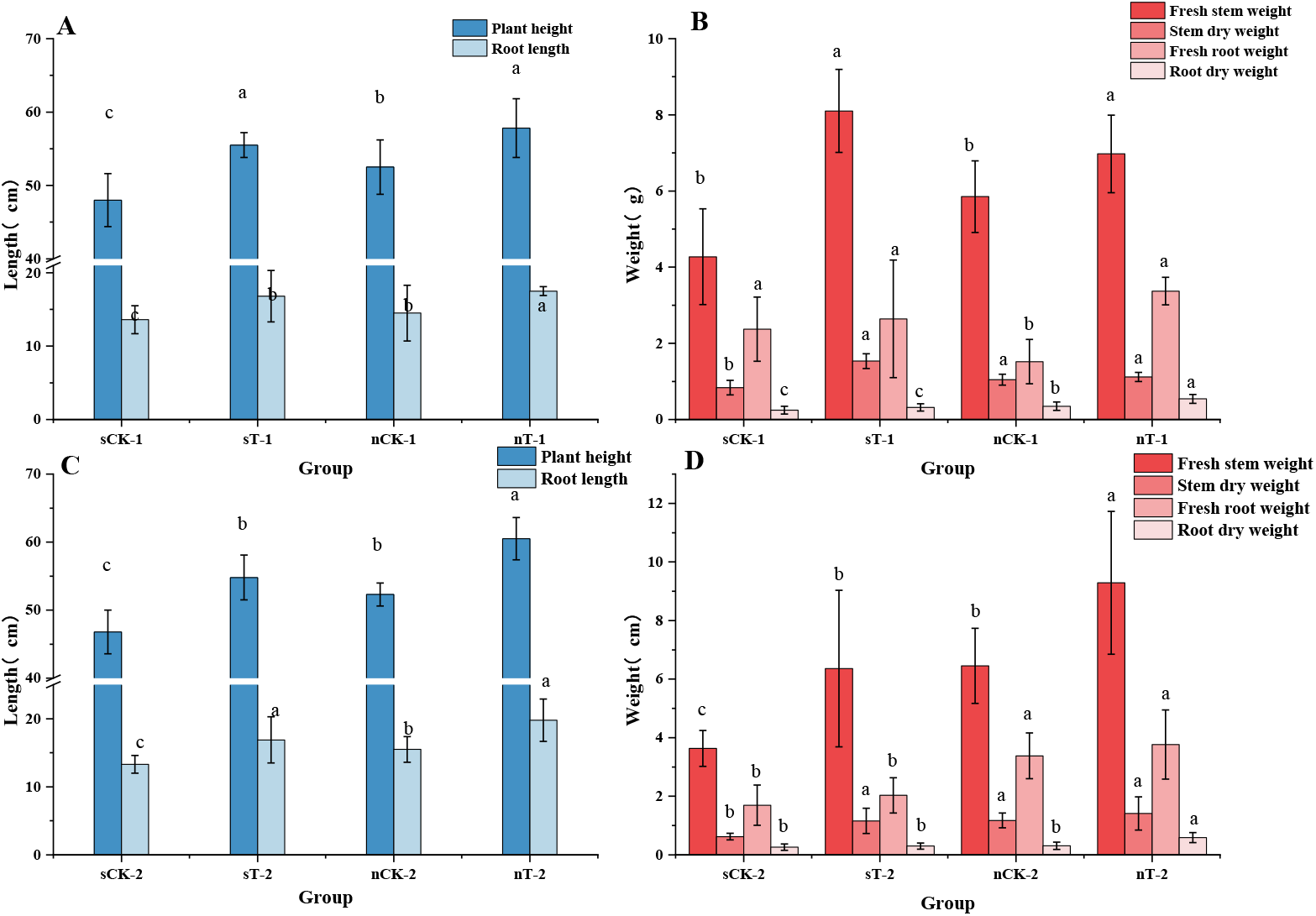
S8 The growth of rice plants in simulated rice fields conditions in 21d. Note that different letters indicate significant differences at the 0.05 level. A : plant height and root lenght of experimental group and control group at recommended dose (375 g·hm^−2^·a^−1^); B: fresh stem weight, stem dry weight, fresh root weight and root dry weight of experimental group and control group at recommended dose (375 g·hm^−2^·a^−1^); C: plant height and root height of experimental group and control group at 2 times recommended dose (750 g·hm^−2^·a^−1^); D: fresh stem weight, stem dry weight, fresh root weight and root dry weight of experimental group and control group at 2 times recommended dose (750 g·hm^−2^·a^−1^).

Since it was discovered that QNC is mainly degraded by microorganisms in nature, many kinds of QNC degrading bacteria had been screened out, but the researchers only studied its degradation characteristics, etc. There were no relevant reports on the bioremediation of QNC-contaminated rice fields by QNC-degrading bacteria, and no reference basis for the practical application of degrading bacteria in the future. In this experiment, the high-efficiency degrading bacteria selected for QNC were used and potted simulated rice fields were used to study the bioremediation of QNC-contaminated rice fields. The test results shown that the QNC-degradation bacteria D could be adapted to the rice field environment and the rice field soil bacteria could degrade QNC cooperatively. The good repair effect can obviously reduce the negative influence of QNC on rice plant physiology, showing a excellent potential broad prospect for application.

## materials and methods

### Materials

QNC standard (98%, Shanghai Yuanye Biological Company); QNC Wettable powder (50%, Jiangsu Kuaida Agrochemical Co., Ltd.); LB liquid medium: 10 g tryptone, 5 g yeast extract, 10 g NaCl, pH 7.0, 1000 mL distilled water; LB solid medium: 10 g tryptone, 5 g yeast extract, 10 g Nacl, 20 g agar, pH 7.0, 1000 mL distilled water; MSM liquid medium: MgSO_4_·7H_2_O 0.2g, CaCl_2_ 0.02g, KH_2_PO_4_ 1.0g, Na_2_HPO_4_ 1.0g, FeCl_3_ 0.05g, NH_4_NO_3_ 1.0g, 1000 mL distilled water; rice paddy soil and rice seeds are all acquired from China Rice Research Institute.

### Screening of QNC-degrading strains

5g paddy soil sample was added to 100mL MSM liquid medium with QNC of 100 mg·L^−1^. After incubating at 30°C and 180 rpm for 7 days, 5 ml of the medium was transferred to new MSM liquid medium with QNC of 200 mg·L^−1^ for cultivation. After 7 days, repeated the transfer until the concentration of QNC in MSM medium reached 500 mg·L^−1^. The final culture medium was gradient diluted and inoculated on LB solid medium. After culturing at 30°C for 48 hours, a single colony was picked and streaked on the LB solid medium to obtain single colonies. Subsequently, different single colonies were selected and cultured in MSM liquid medium with 100 mg·L^−1^ QNC for 21 days, and then QuEChERS-liquid chromatography-tandem mass spectrometry was used to extract QNC and determine the residual concentration of QNC (32) to select the candidate dominant strain.

### Identification of QNC-degrading strains

The selected strain was inoculated on LB solid medium and cultured at 30°C for 48 hours to observe the morphological characteristics of the colony. Gram stain was employed to identify the gram-positive or gram-negative of the strain and the morphology of bacteria was observed under a scanning electron microscope. Physiological and biochemical kits (Qingdao Haibo Biological Co., Ltd.) were used for physiological and biochemical identification. “Berger’ Bacterial Identification Manual” was referred for the identification. After strain D were activated, the BigDye Teminator V3.1 Cycle Sequencing Kit (ABI, USA) was used to extract the total DNA of strain D. 1% agarose gel electrophoresis was used to detect DNA extraction efficiency. A couple of bacterial universal primer (27F: 5’-AGAGTTTGATCCTGGCTCAG-3’ / 1492R: 5’-TACCTTGTTACGACTT-3’) was used. Amplification reaction system was: DNA 1 μL, BigDye 8 μL, primer (3.2 pm·l·L^−1^) 1 μL, sterile deionized water 10 μL. Reaction conditions was as followed: firstly, pre-denaturation at 96 °C for 1 min; secondly, denaturation at 96 °C for 10 s, annealing at 50 °C for 5 s, extension at 60 °C for 4 min, 25 cycles; finally, storage at 4 °C Amplification products were submitted to Shanghai Shangya Biotechnology Co., Ltd. for sequencing. The sequencing results were uploaded to the NCBI database, and the homology comparison was performed by Blast in GenBank (http://www.ncbi.nlm.nih.gov). The phylogenetic analysis was carried out by MEGA7.0 software, and the phylogenetic tree was constructed by the Neighbor-Joining method.

### The degradation characteristics of QNC by strain D

#### Preparation of bacteria suspension

Strain D was inoculated into 100 ml liquid LB medium, incubated at 30°C, 180 rpm for 48 hours, centrifuged at 8000 rpm for 5 minutes, and then poured off the upper layer of LB. The precipitated strain was washed twice with sterile physiological saline solution, and then resuspended the pellet in MSM liquid medium (*OD*_600_=1.0) for later use.

#### Effects of different culture conditions on the degradation rate of QNC

The effects of pH, initial QNC, inoculum amount, yeast extract content, N source and temperature on QNC degradation were investigated. The pH of the culture solution was adjusted to 4, 6, 7, 8, 10 respectively, and culture was performed at 30 °C MSM liquid medium was made up with QNC concentration of 20, 50, 100, 200, and 500 mg·L^−1^, respectively,cultured at 30 °C The initial Inoculation amount of the culture solution was set to 1%, 2%, 5%, 7% and 10%, respectively, cultured at 30 °C Add 0, 0.1%, 0.5%, 1% and 2% of yeast extract to investigate the effect on QNC degradation. Urea, peptone, (NH_4_)_2_SO_4_, NH_4_NO_3_ and NH_4_CL was added individually to cultures at 30 °C to assess the effect of the N source. The medium was cultured at 20 °C 25 °C 30 °C 35 °C and 40 °C to assess the effect of temperature. Samples were taken at 3, 7, 12, 15, 18 and 21 days after inoculation to determine the concentration of QNC and calculate the degradation rate. Three parallels were set for each treatment, and the treatment without inoculation was set as the control.

#### Degradation of QNC by strain D in optimal culture conditions

The MSM liquid medium with a QNC concentration of 50 mg·L^−1^ was prepared, adjust the pH to 6. The N source of MSM medium was (NH_4_)_2_SO_4_, added with 0.1% yeast extract powder, and 7% bacterial suspension. The culture condition of MSM medium was 30 °C and 180 rpm. Samples were taken at 3, 7, 12, 15, 18 and 21 days after inoculation to determine the concentration of QNC and calculate the degradation rate. And the OD(600) value of the bacterial suspension was determined by using an ultraviolet spectrophotometer (UV2600, Shimadzu, Japan). Three parallels were set for each treatment, and the treatment without bacteria was set as the control.

#### Determination of degradation products

Strain D was cultured in optimal conditions for 21d, and samples were taken at 3, 7, 12, 15, 18 and 21 days after inoculation. The QuEChERS method was used to extract QNC and its degradation products. The degradation products of QNC were detected by high performance liquid chromatography tandem quadrupole-time-of-flight mass spectrometer (HPLC-Q-TOF/MS, Agilent, USA). The HPLC conditions were as follows: Xselect HSS T3 column (American waters, 2.1 mm × 150 mm × 3.5 μm); column temperature, 45°C; aqueous solution containing 0.1% formic acid (v/v) (A) and acetonitrile (B) as the flow Phase, gradient elution is 0-5 min, 10%-40% B; 5-11 min, 40%-95% B; 11-15 min, 95% B; 15-15.1 min, 95%-10% B; 15.1-21 min, 10% B; flow rate 300 μL·min^−1^, injection volume is 5 μL. Mass spectrometry conditions: The electrospray source used in the mass spectrometer is ESI, using positive and negative ion full scan modes, the mass scan range m/z 50-1200, and the mass scan rate is 4 spectra·s^−1^. The parameters of positive and negative ion scanning are set as follows: drying gas temperature 350°C drying gas flow rate 8 L·min^−1^, sheath gas temperature 280°C sheath gas flow rate 11 L·min^−1^, atomizing gas pressure 40 psig, fragmentation voltage 130 V, cone The skimmer voltage is 40 V, and the RF voltage is 750 V. Positive ion mode: capillary voltage 4000 V, nozzle voltage 500 V; negative ion mode: capillary voltage 3500 V, nozzle voltage 1000 V.

### The evaluation of strain D on the degradation of QNC and the growth of rice plants in simulated QNC-contaminated rice fields

#### Pre-treatment for pot experiment

Soil: Paddy soil was dried and sieved (2mm), and part of the soil was sterilized; Strains: strain D was inoculated in LB liquid medium, cultured at 30°C, 180 rpm for 48 h, then washed twice with physiological saline, resuspended in steriled water to obtain a bacterial suspension for later use; Pesticide preparation: According to the recommended dosage of QNC and 2 times the recommended dosage, the mother liquor of QNC with an active ingredient of 375 g·hm^−2^·a^−1^ and an effective ingredient of 750 g·hm^−2^·a^−1^ was prepared for use; Rice seeds: the rice seeds were sterilized and soaked to accelerate germination, and cultivated at the two-leaf one-heart stage in an artificial incubator. Rice seedlings of uniform growth were selected for transplanting.

#### Pot experiment

Selected rice seedlings that grow consistently and robust were transplanted into potting buckets (25cm×20cm) with 6 kg of soil in each bucket. After 7 days of transplanting, 8 treatments were carried out respectively: nCK-1: normal soil, sprayed QNC according to the recommended dosage (375 g·hm^−2^·a^−1^). nT-1: normal soil, sprayed QNC according to the recommended dosage, added the bacterial suspension and mixed evenly to make the concentration of 3.38×10^8^ CFU·g^−1^; nCK-2: normal soil, applied twice of the recommended dosage (750 g·hm^−2^·a^−1^). nT-2: normal soil, apply 2 times of the recommended dose, and added the bacterial suspension and mixed evenly to make the concentration of 3.38×10^8^ CFU·g^−1^; sCK-1: sterilized soil, applied the recommended dose. sT-1: sterilized soil, applied QNC according to the recommended dose, added the bacterial suspension and mixed evenly to make the concentration of 3.38×10^8^ CFU·g^−1^; sCK-2: sterilized soil, applied twice of the recommended dose. sT-2: sterilized soil, apply 2 times of the recommended dose, added the bacterial suspension and mixed evenly to make the concentration of 3.38×10^8^ CFU·g^−1^. Each treatment was repeated for three times. After application, random multi-point sampling was used at 0, 3, 7, 14, 21, and 28d to collect rice plants and soil samples. Soil samples were employed to detect QNC residues. The plant height, root length, stem dry weight, stem fresh weight, root dry weight and root fresh weight of rice plant samples collected at 7d and 21d were measured respectively.

## Data availability

The complete 16S rDNA sequence of Cellulosimicrobium sp D have been deposited in GenBank under the accession no. NR_119095.

## Acknowledgements

This work was founded by the Special Fund for the Construction of Modern Agricultural Industrial Technology System (CARS-01-47), Collaborative Innovation Task of the Science and Technology Innovation Project of the Chinese Academy of Agricultural Sciences: Product Creation and Demonstration Application of Quick Test Product Quality and Safety of Agricultural Products (CAAS-XTCX2019024).

